# Temporal and Spatial Characterization of CUL3^KLHL20^-driven Targeted Degradation of BET family, BRD Proteins by the Macrocycle-based Degrader BTR2004

**DOI:** 10.1101/2024.12.07.627262

**Authors:** Phoebe H. Fechtmeyer, Johannes T.-H. Yeh

## Abstract

Targeted protein degradation (TPD) is a promising new therapeutic modality that leverages the endogenous cellular protein degradation machinery of the ubiquitin-proteasome system (UPS) to degrade selected proteins. Recently, we developed a synthetic macrocycle ligand to recruit CUL3^KLHL20^ E3 ligase for TPD. Using this KLHL20 ligand, we constructed the PROTAC BTR2004, which demonstrated potent degradation of BET family proteins BRD 2, 3, and 4. As the TPD field expands, it is important to understand the cellular and biochemical properties of all utilized E3 ligases. Herein we report the temporal and spatial processes of BTR2004-facilitated BET family protein degradation by KLHL20: The target protein degradation kinetics, BTR2004 intracellular activity half-life, and the onset of BTR2004 cell permeabilization. Employing proximity ligation and confocal microscopy techniques, we also illustrate the subcellular location of the ternary complex assembly upon BTR2004 treatment. These characterizations provide further insight into the processes that govern TPD and features that could be incorporated when designing future PROTAC molecules.

## Introduction

The past few years have seen immense interests in the Targeted Protein Degradation (TPD) approach to drug discovery and experimental therapeutics development^1^. Although numerous protein degraders have been generated to target a wide variety of proteins, most degrader molecules engage CRBN and VHL E3 ligases. As therapeutic applications continue to grow, expanding the E3 ligase landscape to broaden TPD potential has been suggested in the field^2^. In 2022, we reported the functional validation of the E3 ligase CUL3^KLHL20^, adding KLHL20 as a new utilizable E3 ligase for TPD^3^.

KLHL20 is one of more than 60 BTB-Kelch family adaptor proteins that belong to the CUL3 superfamily. First characterized as an oncogene^4,5^, KLHL20 is ubiquitously expressed in many cancer cells, making it an ideal E3 ligase to execute TPD in cancer treatment. In our functional validation of KLHL20 we designed a small macrocyclic peptide ligand, BTR2000, which binds to the degron binding pocket in the Kelch domain of KLHL20^3,6^. Strikingly, proteomics analysis demonstrated that BTR2000 exhibits a high specificity to KLHL20, even though it contains only three amino acid residues for KLHL20 binding^3^. BTR2000 thus can serve as a selective KLHL20-recruiting ligand for PROTAC molecule expansion, akin to the thalidomide analogs to CRBN-based PROTACs^7^ or the VH032 analogs to VHL-based PROTACs^8^. As an example, we made BET-family BRD protein degraders by conjugating the Bromodomain inhibitor JQ1 to BTR2000 via a linker. From the small library of compounds, BTR2004 was identified as a potent BRD2, BRD3 and BRD4 degrader^3^.

While CRBN- and VHL-driven TPD has been widely documented^7^ and characterized^9^, KLHL20-driven TPD by BTR2000-derived PROTACs is still new to the field. BTR2004, as a macrocyclic peptide-small molecule linked heterobifunctional PROTAC (Figure 1A), represents a novel class of BRD protein degrader. To better understand the bioactive properties of this new type of PROTAC, we aim to further characterize KLHL20-driven TPD using BTR2004. Additionally, previous studies have characterized the kinetics of degradation^9-11^, but few have evaluated the kinetics of PROTAC cell entry or the subcellular localization of ternary complex formation. In this study, we performed temporal and spatial characterizations on BTR2004-facilitated BRD protein degradation such as degradation speed, duration of efficacy, compound cellular entry, and subcellular localization of ternary complex formation.

**Figure 1.**
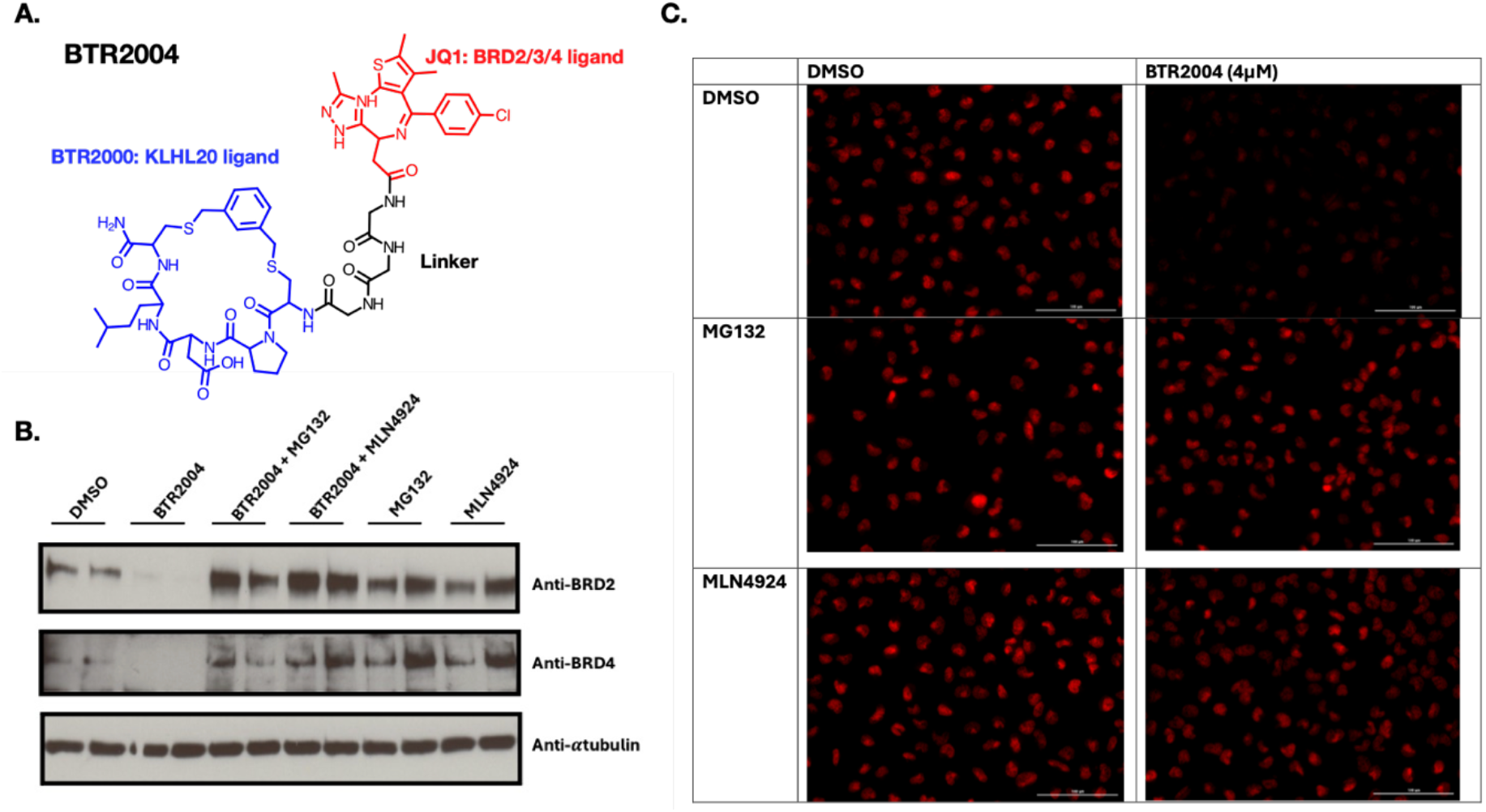
BTR2004-facilitated targeted protein degradation is ubiquitin-proteasomal pathway dependent. **(A)** Chemical structure of BET family protein PROTAC: BTR2004. **(B)** Western blot of BRD2/4 degradation in U20S T-REx cells treated with BTR2004 (8μM), MG132 (10μM), MLN4924 (3μM), and DMSO. MG132 and MLN4924 prevented BTR2004 facilitated BRD2 and BRD4 degradation. **(C)** Immunofluorescent staining of BRD4 (Red) in U20S T-REx cells treated with BTR2004 or DMSO with or without MG132 or MLN4924 co-treatment.

## Results

### BTR2004 facilitates target protein degradation via the Ubiquitin-Proteasome System

Our previous study^3^ validated KLHL20 as a potent E3 ligase suitable for TPD. Although the Ubiquitin-Proteasome System (UPS) is the only known pathway for CUL3^KLHL20^ driven protein degradation, we wanted to confirm that BTR2004-facilitated BRD protein degradation was indeed UPS pathway-dependent. The Western blot results exemplified that BRD2 and BRD4 degradation by BTR2004 was prevented when cells were treated with the proteasome inhibitor MG132 or the Nedd8-activating enzyme inhibitor MLN4924 (Figure 1B). A similar finding was observed by immunofluorescent (IF) imaging of BRD4 (Figure 1C). These results confirmed that BTR2004 mediated BRD protein degradation is UPS pathway dependent.

### BTR2004 degradation kinetics

Among the three BET family BRD proteins, BTR2004 has higher activity towards BRD2^3^. Therefore, in this study, the characterization focused on BRD2 protein level as the readout. In the kinetics measurements of BTR2004-facilitated BRD2 degradation, ≥ 2 μM of BTR2004 was chosen to assure the maximum BRD2 degradation (Figure 2A). Overall, 4 μM of BTR2004 efficiently eradicated BRD2 protein in ∼ 2 hours, however, no observable degradation occurred until approximately 40 minutes after compound introduction (Figure 2B). To delineate the degradation kinetics, BRD2 protein levels were plotted at different BTR2004 incubation durations, using concentrations of 2 μM and 4 μM BTR2004. The degradation plot followed a sigmoidal shape, and both curves could be fit using a sigmoidal equation (Figure 2C). As anticipated, the protein degradation rate was compound concentration-dependent: 4 μM of BTR2004 treatment exhibited a faster degradation rate and achieved maximum degradation sooner than 2 μM treatment. Interestingly, for both 2 μM and 4 μM BTR2004, degradation initiation is observed around the same time and a similar maximum degradation value is reached.

**Figure 2.**
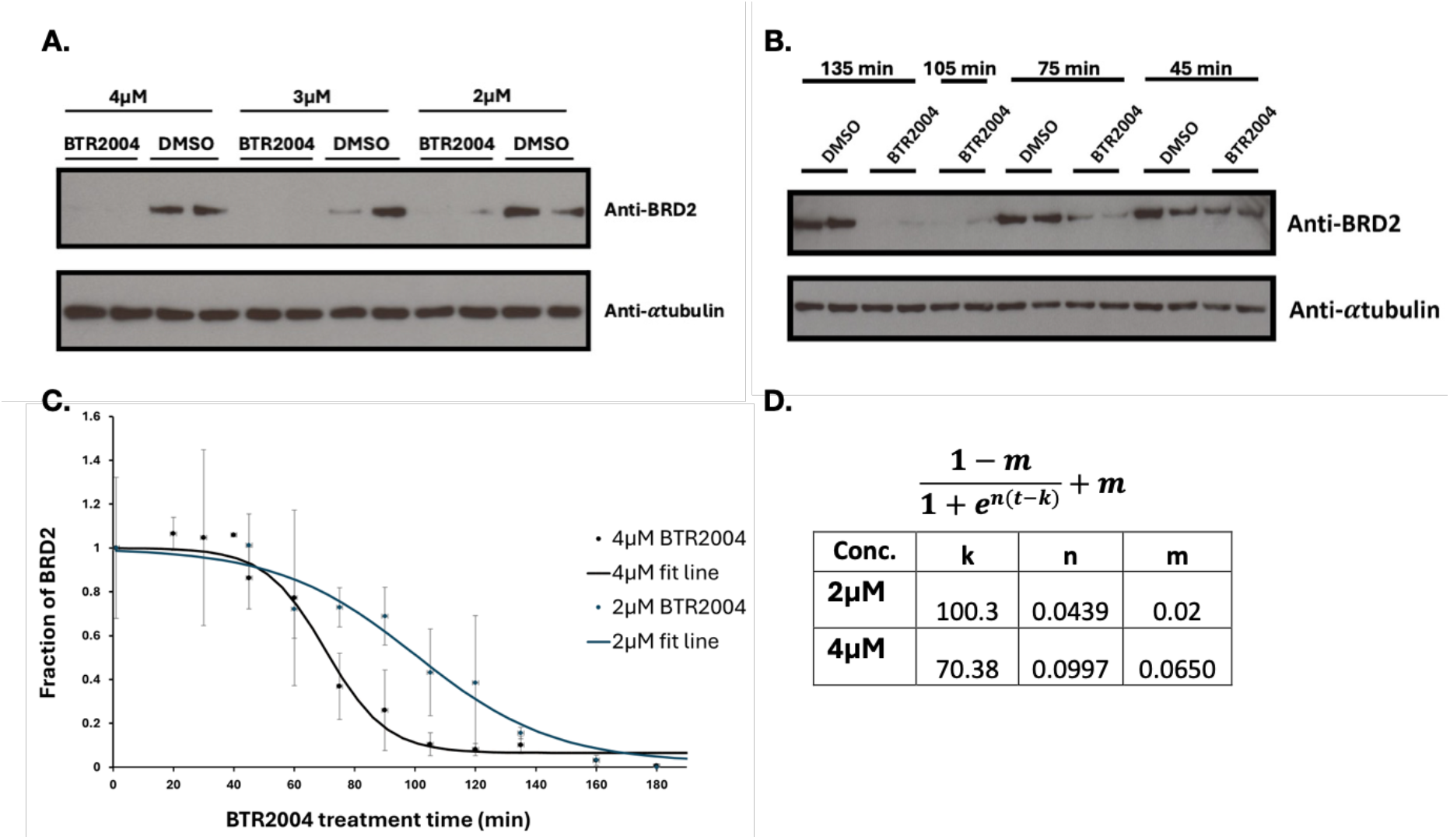
BTR2004 degradation kinetics. **(A)** Western blot of BRD2 degradation with different BTR2004 concentrations. **(B)** BRD2 degradation by 4μM BTR2004 with different treatment time. **(C)** Quantitative BRD2 degradation time course with 2 μM and 4 μM BTR2004 treatment showed different kinetics. **(D)** Figure 2C curve fitting parameters: The constant “n,” is proportional to the slope of the line and therefore related to BRD2 degradation rate (normalized protein/min). The maximum rate of degradation is equal to n(1-m)/4. The constant “k” is the time (minutes) at which protein is being maximally degraded and refers to the time at which half of the protein has been attenuated. The constant “m” is equal to remaining protein % at maximum degradation.

By fitting sigmoidal curves, the constants calculated can be used to directly compare the kinetic behavior at different concentrations (Figure 2D). The constant “m” is the minimum, therefore equal to the fraction of protein remaining at maximum degradation. The constant “k” is the time of the inflection point, where degradation rate is maximum, and halfway to maximum degradation. The constant “n” is proportional to the slope and therefore degradation rate (normalized protein/min). The maximum rate of degradation was calculated by the equation: 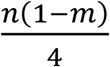 It can be observed that for 2 μM treatment n= 0.044 (∼1.1% BRD2 degradation/minute at the maximum degradation speed) and k= 100 (100 minutes for half of maximum BRD2 degradation) while with 4 μM treatment n= 0.0999 (∼2.3% BRD2 degradation/minute at the maximum degradation speed) and k= 70 (70 minutes for half of maximum BRD2 degradation).

### Degradability persistence and bioactivity half-life of BTR2004

Besides protein degradation rate, which is maintained by the constant presence of degrader at the equilibrium state, another important aspect of a degrader’s efficacy is its persistence in the cell. To test the persistence of BTR2004’s activity in the cell, we performed a compound “washout” experiment. In this setting, U2OS T-REx cells were incubated with media containing 4 μM BTR2004 for 2 hours to reach the maximum degradation of BRD2, then the cells were washed twice and replenished with the culture medium (without BTR2004) for a continued culture period of variable times (Figure 3A). In this “washout” experiment, sustained BRD2 degradation can only be carried out by the BTR2004 compounds that entered the cell during the initial 2-hour compound incubation time. Subsequently, the intracellular bioactivity persistence of BTR2004 can be determined based on the duration that BRD2 remained degraded. BRD2 protein remained maximumly degraded for 2 hours after BTR2004 washout. After that, the protein level gradually increased and fully recovered around 6∼8 hours after BTR2004 washout (Figure 3B). By contrast, BRD2 protein levels remain maximally degraded for at least 25 hours under constant incubation with 4 μM BTR2004 media (Figure 3C). The findings indicated that the full cellular activity of BTR2004 can last for at least 2 hours and then would gradually fade away between 2∼8 hours without continued compound replenishment.

**Figure 3.**
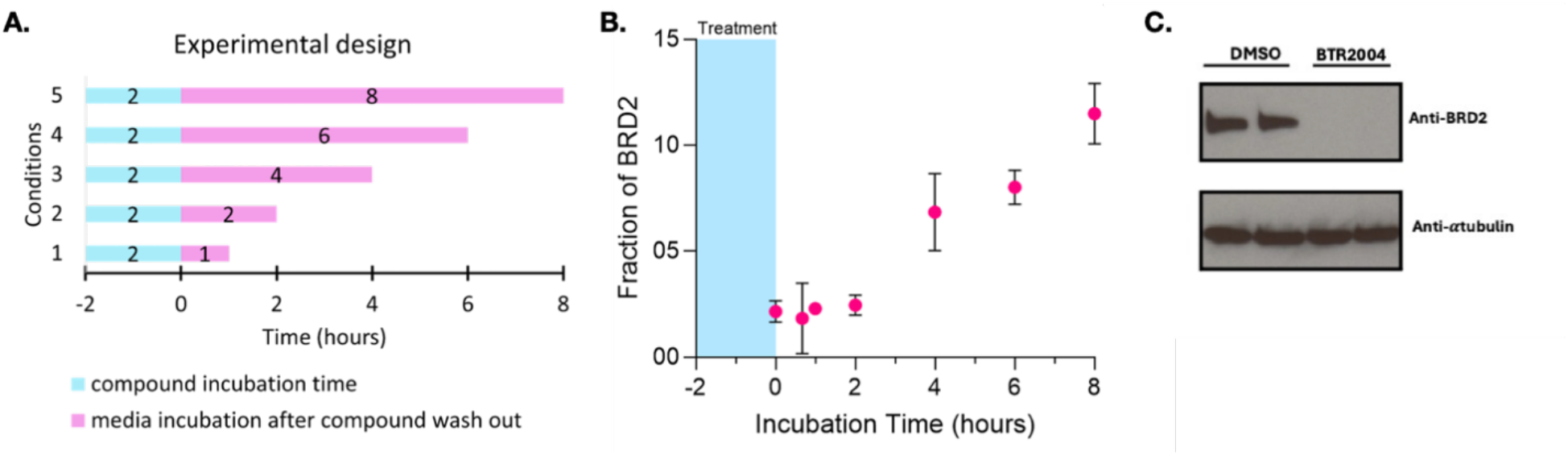
Degradability persistence and bioactivity half-life of BTR2004. **(A)** Schematic diagram of compound washout experiment design. U2OS T-REx cells were first incubated with 4 μM BTR2004 for 2 hours (highlighted in blue) followed by compound washout and continued incubation for varying periods of time (highlighted in pink) in compound-free culture medium. **(B)** Cellular BRD2 protein recovery level over time after BTR2004 washout. Protein level measured by western blot quantification. **(C)** BRD2 protein remained degraded for 25 hours with constant BTR2004 incubation.

### Cell permeability of BTR2004

From the cellular BRD2 degradation time-course curve (4 μM BTR2004), we noticed a lag phase in the first ∼60 minutes after BTR2004 treatment. This lag phase could be attributed to the time required for BTR2004 cell entry, ternary complex formation, BRD2 polyubiquitination and proteasomal degradation. However, we speculated that the time required for sufficient BTR2004 cell entry was the primary limiting step accounting for the time lag. To elucidate the earliest timepoint at which BTR2004 reached the intracellular concentration sufficient for catalyzing BRD2 degradation, we set up a modified washout experiment (Figure 4A) based on the facts that BTR2004 achieved maximum BRD2 degradation at ∼2 hours and the intracellular bioactivity can persist for at least 2 hours. We therefore let the cells incubate with 4 μM BTR2004 for varying times less than 2 hours (ranging from 5∼65 minutes), then washed out BTR2004 and allowed the cells to continue incubating in compound-free medium for the remaining time (to reach a total 135-minutes incubation time counting from BTR2004 introduction). In this experiment, we demonstrated that with a total 135-minutes culture duration, even 65 minutes of BTR2004 exposure could still reach degradation level comparable to the full course BTR2004 exposure (i.e., 135 minutes BTR2004 incubation) (Figure 4B). This phenomenon indicated that 65 minutes of BTR2004 exposure was sufficient to drive an autonomous TPD process in the exogenous compound-free culture condition, as long as the cellular process are not prematurely stopped (i.e., < 135 min). Along the same line, as BTR2004 exposure time shortened to 40 minutes ∼50% of BRD2 protein reduction was observed. Strikingly, observable signs of BRD2 degradation could be seen with as little as 20 minutes of BTR2004 treatment (Figure 4B). Based on our findings collectively, we concluded this working model: Although initial degradation of BRD2 did not occur until ∼ 60 minutes after BTR2004 treatment, the compound had entered the cells at a much earlier time point (< 20 mins) and started to accumulate. Then around the 40th-minute, the cellular concentration of BTR2004 would reach the DC_50_ level, while between 40∼60 minutes the accumulated cellular BTR2004 concentration was sufficient for maximum level of BRD2 degradation (Figure 4B). Our characterization thus suggested that the first hour lag phase was critical to allow sufficient cellular accumulation of BTR2004.

**Figure 4.**
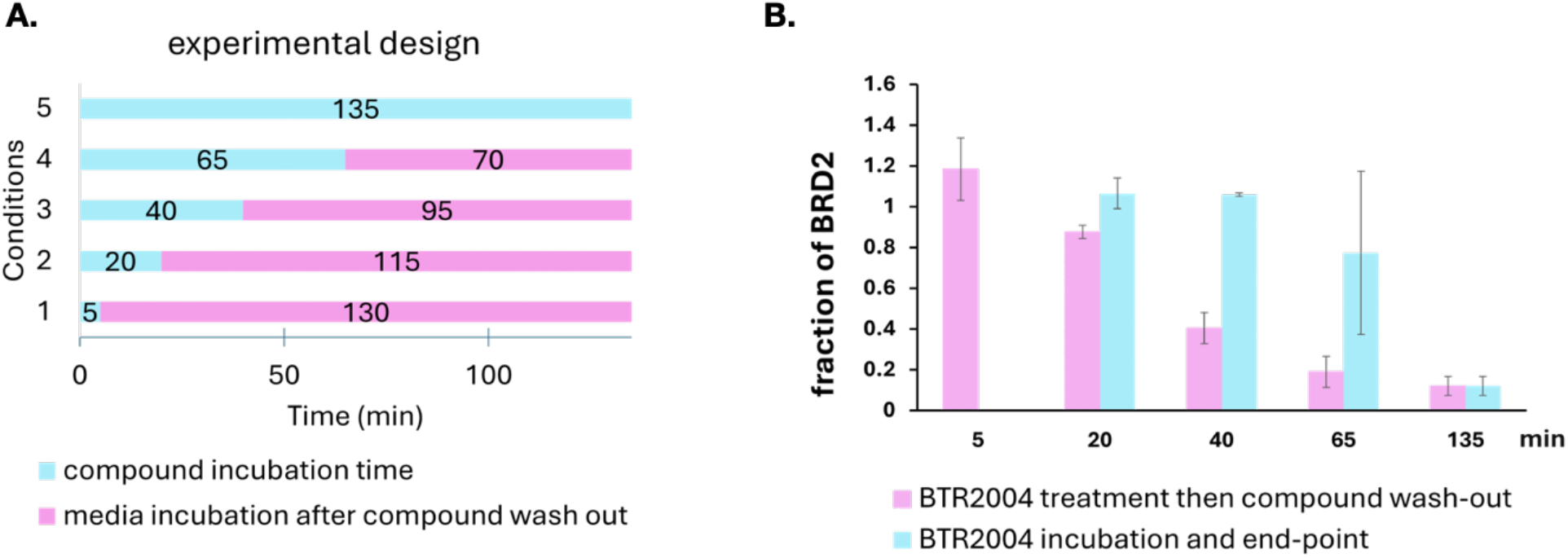
BTR2004 bioactivity onset. **(A)** Schematic diagram of modified BTR2004 washout experiment design. U2OS T-REx cells were first incubated with BTR2004 for varied amounts of time (highlighted in blue) followed by compound washout and continued incubation in compound-free culture medium until 135 minutes total incubation endpoint (highlighted in pink). **(B)** BRD2 protein level at 135 minutes total incubation endpoint (pink) vs. BRD2 protein level at the end of BTR2004 treatment (blue).

### Subcellular localization of ternary complex formation

We next were curious to know where BTR2004 facilitated protein ternary complex inside the cells. The known subcellular localization of BET family proteins is strictly nuclear^12^. However, we found a vast majority of the KLHL20 localized in the cytosol and only hints of nuclear localization (Figure 5A). Due to the lack of antibodies suitable for Immunofluorescence (IF) staining with KLHL20, we had to introduce an HA-tagged KLHL20 (HA-KLHL20) construct into the U2OS T-REx cell line under the tetracycline-inducible expression control (Tet-on). Using the titratable Tet-On system, we tuned the tetracycline concentration to make the HA-KLHL20 expression level just detectable by IF imaging without overly expressing the protein. This allowed us to visualize the localization of HA-KLHL20 by an anti-HA antibody.

**Figure 5.**
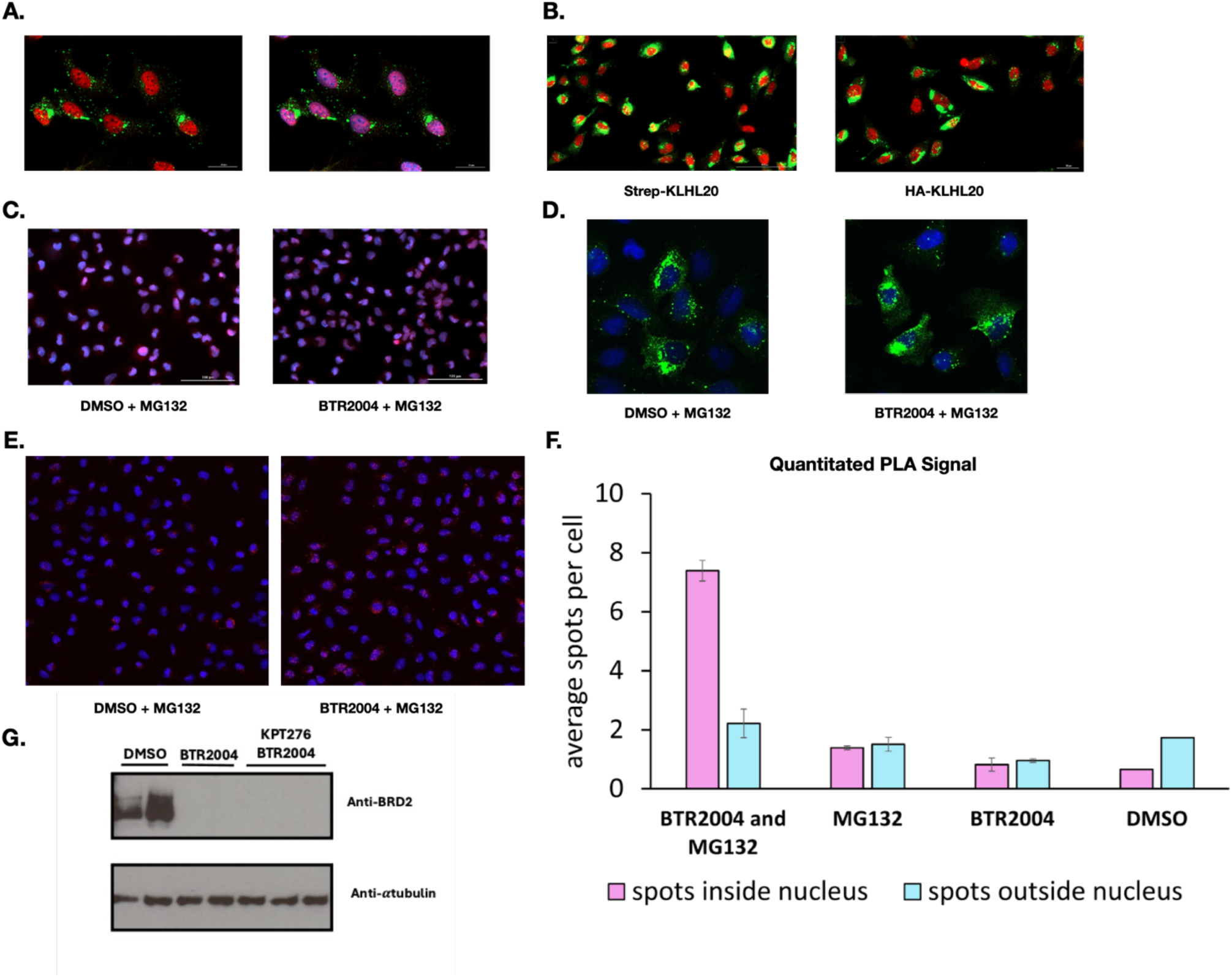
Subcellular localization of ternary complex formation. **(A)** BRD4 (red) is exclusively localized to the nucleus whereas most HA-KLHL20 (green) is cytosolic. (*Left*) Merged image of BRD4 (red) and HA-KLHL20 (green) channels. (*Right*) Merged image with DAPI staining. (**B)** Strep-KLHL20 and HA -KLHL20 (green) showed similar localization patterns. **(C)** BTR2004 did not change BRD4 (red) subcellular localization. DAPI nuclear staining in blue. **(D)** BTR2004 did not change KLHL20 (green) subcellular localization. DAPI nuclear staining in blue. **(E)** BTR2004 mediated ternary complex formation revealed by Proximity Ligation Assay (PLA) using confocal microscopy imaging. (**F)** Quantified PLA signals (using Imaris program) showed ternary complex primarily formed in the nucleus. **(G)** Exportin 1 inhibitor KPT276 did not prevent BRD2 degradation, suggesting nuclear proteasomal degradation of the nuclear formed ternary complex.

To exclude the possibility that the HA tag was influencing KLHL20’s localization, we also constructed a Strep tagged KLHL20 (Strep-KLHL20) Tet-on cell line. Both HA-KLHL20 and Strep-KLHL20 showed consistent cytosolic localization patterns, indicating the epitope tags had little artificial influence on KLHL20 subcellular localization (Figure 5B). It is also worth noting that we chose HA and Strep tags because they do not contain any Lysine residues that could potentially be ubiquitinated and artificially alter the tagged KLHL20’s protein stability (e.g., leading to self-polyubiquitination of KLHL20).

Interestingly, the introduction of BTR2004 did not affect the subcellular localization of either KLHL20 or BRD2 (Figure 5C-D). This finding therefore posed the question: where does the KLHL20-BTR2004-BRD protein ternary complex form upon BTR2004 engagement?

To further detect the localization of the KLHL20-BTR2004-BRD ternary complex, we employed the Proximity Ligation Assay (PLA) to label the ternary complex for cell imaging. Since PLA requires the two primary antibodies derived from two different animal species for simultaneous secondary antibody labeling (e.g., anti-mouse & anti-rabbit secondary antibody), we had to employ a mouse anti-human BRD4 antibody for its better performance in IF staining with fixed cells. We thus focused on visualizing the CUL3^KLHL20^-BTR2004-BRD4 ternary complex by PLA. Upon treatment with BTR2004 (and MG132 to stabilize BRD4), the PLA signals were significantly increased as compared to the DMSO and MG132 alone. This confirmed the PROTAC-dependent ternary complex formation. Additionally, most of the PLA signals were nuclear (Figure 5E). We quantified the signal intensity and localization using the Imaris 3D image analysis software (Figure 5F). Since BTR2004 treatment did not affect KLHL20 or BRD4’s cyto-nuclear distribution, this observation indicated that the pre-existing nuclear KLHL20 pool was the primary source to engage ternary complex formation. Despite its low abundance, this pre-existing nuclear KLHL20 pool was sufficient to eradicate the nuclear target proteins. Additionally, adding the Exportin1 inhibitor KPT276 had no effect on the BRD protein degradation (Figure 5G), therefore extending the conclusion that not only was the ternary complex mostly formed in the nucleus, but also that proteasome degradation occurs primarily in the nucleus.

## Discussion

TPD strategy has recently been extensively embraced by the drug discovery community. Featuring the eradication of the target protein in a catalytic manner, PROTACs provide advantages that can compensate for the drawbacks experienced with the conventional inhibitors. While currently, nearly all the clinical stage PROTACs are CUL4^CRBN^ E3 ligase-based, investigations on exploiting new E3 ligases for TPD have begun to grow. Among the ∼ 600 E3 ligases, thus far only a small number of new E3 ligases have been pharmacologically verified (by using a synthesized PROTAC compound to demonstrate degradability) for TPD activity. Besides the well-established CUL4^CRBN^ and CUL2^VHL^, other examples include cIAP^13^, MDM2^14^, RNF4^15^, RNF114^16^, KEAP1, FEM1B^17^, KLHL20 and FBXO22^18^.

For designing a PROTAC strategy, several factors need to be considered, including the choice of the ligase. Since not all E3 ligase are created equal, the enzymatic activities and subcellular localization will significantly affect the kinetics and persistence of target protein degradation. In addition to the intrinsic biochemical and subcellular properties, the physical chemical properties of the PROTAC molecule itself play a critical role in pharmacokinetics and pharmacodynamics.

Here, we have characterized the temporal and spatial activities of the CUL3^KLHL20^-driven BRD protein PROTAC BTR2004. As shown with other PROTACs^9^, BTR2004 concentration can affect the degradation rate, particularly within the short-term timeframe. Furthermore, such a pharmacological behavior is not visible by a single timepoint readout when maximum degradation phase has been reached. For example, both 2 μM and 4 μM treatments of BTR2004 will eventually reach a similar maximum degradation albeit 4 μM BTR2004 can provide a faster acute degradation of BRD2. This would be a useful pharmacokinetic parameter should acute TPD be necessary. Regarding the kinetics, while other studies have fit an exponential decay curve to their degradation data^9,10^, we instead found that a sigmoidal curve fits the best with our plot. Additionally, it encompasses the lag phase at the beginning of degradation, caused by the time required for compound to accumulate in the cell and for UPS machinery to initiate degradation. This difference could arise from using different BRD2 degradation assay format (Western blot based vs. Luciferase based) and the chemical properties of degrader cell entry onset.

Since the conceptualization of TPD, concerns on the feasibility of PROTACs to get to the clinic have been raised because most of the PROTACs were predicted to contradict with the well-accepted Lipinski Rule-of-five^19^ - with the most notable contradiction being their cell permeability with a molecular weight (MW.) exceeding the cut-off size of MW. 500. Surprisingly, numerous examples have proven that heterobifunctional PROTAC molecules do not need to follow the Lipinski rules, leading to the new definition of drug-like property “beyond the Rule-of-five” (bRo5)^20,21^. As BTR2004 represents a new class of heterobifunctional degrader featuring a small cyclic peptide moiety linked with the small molecule JQ1 moiety, not much has been studied about the cellular permeability on these types of TPD molecules. In this study we devised the time-course washout experiment as bioactivity-based readout to demonstrate that BTR2004 entered the cells shortly (< 20 minutes) after compound introduction. Although it is not confirmed what the route of BTR2004’s cell entry is, we speculate that BTR2004 entered the cells in a similar fashion to cell permeable macrocyclic peptides such as Cyclosporin^22^.

Enhancing cell permeability and oral bioavailability are highly desired for macrocyclic peptide drug development. Several remarkable milestones have been made in recent years^23,24^. It will be interesting to apply chemical modification strategies (e.g., amide-to-ester substitution^25,26^, Nmethylation^26^, hydrogen bonding optimization^27^ etc.) to develop oral-bioavailable version of BTR2004 in the near future to gain new insights on engineering this type of PROTAC.

Regarding the subcellular location of E3-neosubstrate complex formation, the intuitive assumption would be that E3 ligase-target protein co-localization is necessary. However, recent studies have offered some arguments that pre-existing co-localization may not be a prerequisite determinant for the degradability of a E3 ligase-neo substrate pair. The hetero-functional molecule itself can facilitate the delocalization of the proteins to achieve proximity-based targeted effects. For example, Gibson *et al*.^28^ showed that a hetero-functional molecule NICE-01 (designed to have both Bromodomain and FKBP12^F36V^ binding moieties) can induce the nuclear import of the cytosolic FKBP ^F36V^ to form ternary complex in the nucleus. Interestingly, in the case of BTR2004, neither KLHL20 nor BRD proteins were cyto-nuclear translocated by the PROTAC. Rather, our results indicated that the existing nuclear pool of KLHL20 was the source for BTR2004-facilitated ubiquitination complex. This finding supports the notion that the target protein and E3 ligase in the same subcellular compartment is favored^29^. With limited examples, currently it is hard to predict the protein translocation potential by any hetero-functional molecules until more studies can be carried out. Systemic characterizations on a variety of protein pairs with different types of hetero-functional molecules can enhance our knowledge to decipher the biological rules of compound-mobilized protein translocation. As proximity-based drugs are heralding a new era of drug discovery innovation^30^, characterizing the spatial and temporal kinetics of PROTAC molecules with the associated E3 ligases and target proteins will significantly help the TPD field to develop better protein modulating TPD molecules.

## Materials and Methods

### Cell lines

U2OS T-REx cells were cultured in RPMI 1640 media with 10% FBS. Tet-on HA-KLHL20 and Strep-KLHL20 stable cell lines were generated by co-transfecting U2OS T-REx cells with pcDNA5 FRT/TO HA-KLHL20 or Strep-KLHL20 with pOG44 plasmid per manufacturer’s user manual (invitrogen). The stable cell lines were selected for using 50 μg/ml hygromycin media.

### Immunofluorescent (IF) staining

Cells were plated in a 96 well plate and treated for 3 hours then fixed with 4% Paraformaldehyde (PFA.) IF staining was done with the primary antibodies mouse-anti-BRD4 (E4X7E) and rabbitanti-HA (C29F4) from Cell Signaling Technologies and secondary antibodies AlexaFluor647 conjugated goat-anti-mouse (cat. # 126932) and AlexaFluor488 conjugated goat-anti-rabbit (cat. # 137069) from Jackson Immuno-Research. The cells were imaged on the Cytation 5 plate reader, the Leica SP8 confocal microscope or the Zeiss LSM780 confocal microscope.

### Proximity Ligation Assay

U2OS T-REx HA-KLHL20 cells were plated in a 96 well plate, pretreated with MG132 for 1 hour then treated with 12 μM of BTR2004 for 3 hours. Although 4 μM is sufficient for maximum degradation, 12 μM was used to maximize the numbers visualizable ternary protein complex. After treatment, the cells were fixed with 4% PFA. The Proximity Ligation Assay (PLA) kit was purchased from Navinci (NaveniFlex Cell MR Atto647N). PLA procedure was carried out per manufacturer’s protocol, using a reaction volume of 35 μL per well: After fixation, cells were blocked and incubated with primary antibodies mouse-anti-BRD4 (E4X7E) and rabbit-anti-HA (C29F4) overnight. The next day, the cells were washed and incubated with the provided secondary antibodies, then incubated separately in two reaction mixes. Final imaging was acquired by the Zeiss LSM780 confocal microscope.

## Acknowledgments

We thank Dr. Erika Wee at CSHL Microscopy Shared Resource for assistacne with confocal microscopy and image analysis; Brian Farrell and Carmelita Bautista for assistance with research reagents; CSHL Antibody and Phage Display Shared Resource for providing anti-α tubulin (clone DM1A) monoclonal antibody; Drs. Yousef Al-Abed, Mingzhu He and Kai F. Cheng at Feinstein Institute for Medical Research for instrument access; and Cameron Martinez for assistance with manuscript preparation. This work was supported in part by the CSHL Shared Resource funded by the Cancer Center Support 5P30CA045508, and the Cold Spring Harbor Laboratory and Northwell Health Affiliation.

## Notes

### Competing Interest Statement

The authors have declared no competing interest.

## References

1 Tsai, J. M., Nowak, R. P., Ebert, B. L. & Fischer, E. S. Targeted protein degradation: from mechanisms to clinic. Nat Rev Mol Cell Biol 25, 740–757, doi:10.1038/s41580-024-00729-9 (2024).

2 Kannt, A. & Dikic, I. Expanding the arsenal of E3 ubiquitin ligases for proximity-induced protein degradation. Cell Chem Biol 28, 1014–1031, doi:10.1016/j.chembiol.2021.04.007 (2021).

3 Farrell, B. M., Gerth, F., Yang, C. R. & Yeh, J. T. A synthetic KLHL20 ligand to validate CUL3(KLHL20) as a potent E3 ligase for targeted protein degradation. Genes Dev 36, 1031–1042, doi:10.1101/gad.349717.122 (2022).

4 Yuan, W. C. et al. A Cullin3-KLHL20 Ubiquitin ligase-dependent pathway targets PML to potentiate HIF-1 signaling and prostate cancer progression. Cancer Cell 20, 214–228, doi:10.1016/j.ccr.2011.07.008 (2011).

5 Lee, Y. R. et al. The Cullin 3 substrate adaptor KLHL20 mediates DAPK ubiquitination to control interferon responses. EMBO J 29, 1748–1761, doi:10.1038/emboj.2010.62 (2010).

6 Chen, Z., Picaud, S., Filippakopoulos, P., D’Angiolella, V. & Bullock, A. N. Structural Basis for Recruitment of DAPK1 to the KLHL20 E3 Ligase. Structure 27, 1395–1404 e1394, doi:10.1016/j.str.2019.06.005 (2019).

7 Bekes, M., Langley, D. R. & Crews, C. M. PROTAC targeted protein degraders: the past is prologue. Nat Rev Drug Discov, doi:10.1038/s41573-021-00371-6 (2022).

8 Zengerle, M., Chan, K. H. & Ciulli, A. Selective Small Molecule Induced Degradation of the BET Bromodomain Protein BRD4. ACS Chem Biol 10, 1770–1777, doi:10.1021/acschembio.5b00216 (2015).

9 Riching, K. M. et al. Quantitative Live-Cell Kinetic Degradation and Mechanistic Profiling of PROTAC Mode of Action. ACS Chem Biol 13, 2758–2770, doi:10.1021/acschembio.8b00692 (2018).

10 Riching, K. M., Caine, E. A., Urh, M. & Daniels, D. L. The importance of cellular degradation kinetics for understanding mechanisms in targeted protein degradation. Chem Soc Rev 51, 6210–6221, doi:10.1039/d2cs00339b (2022).

11 Schwalm, M. P. et al. Tracking the PROTAC degradation pathway in living cells highlights the importance of ternary complex measurement for PROTAC optimization. Cell Chem Biol 30, 753–765 e758, doi:10.1016/j.chembiol.2023.06.002 (2023).

12 Hsu, S. C. et al. The BET Protein BRD2 Cooperates with CTCF to Enforce Transcriptional and Architectural Boundaries. Mol Cell 66, 102–116 e107, doi:10.1016/j.molcel.2017.02.027 (2017).

13 Schiemer, J. et al. Snapshots and ensembles of BTK and cIAP1 protein degrader ternary complexes. Nat Chem Biol 17, 152–160, doi:10.1038/s41589-020-00686-2 (2021).

14 Hines, J., Lartigue, S., Dong, H., Qian, Y. & Crews, C. M. MDM2-Recruiting PROTAC Oiers Superior, Synergistic Antiproliferative Activity via Simultaneous Degradation of BRD4 and Stabilization of p53. Cancer Res 79, 251–262, doi:10.1158/0008-5472.CAN-18-2918 (2019).

15 Ward, C. C. et al. Covalent Ligand Screening Uncovers a RNF4 E3 Ligase Recruiter for Targeted Protein Degradation Applications. ACS Chem Biol 14, 2430–2440, doi:10.1021/acschembio.8b01083 (2019).

16 Luo, M. et al. Chemoproteomics-enabled discovery of covalent RNF114-based degraders that mimic natural product function. Cell Chem Biol 28, 559–566 e515, doi:10.1016/j.chembiol.2021.01.005 (2021).

17 Henning, N. J. et al. Discovery of a Covalent FEM1B Recruiter for Targeted Protein Degradation Applications. J Am Chem Soc 144, 701–708, doi:10.1021/jacs.1c03980 (2022).

18 Basu, A. A. et al. A CRISPR activation screen identifies FBXO22 supporting targeted protein degradation. Nat Chem Biol 20, 1608–1616, doi:10.1038/s41589-024-01655-9 (2024).

19 Lipinski, C.A. Lead- and drug-like compounds: the rule-of-five revolution. Drug Discov Today Technol 1, 337–341, doi:10.1016/j.ddtec.2004.11.007 (2004).

20 Lipinski, C. A. Rule of five in 2015 and beyond: Target and ligand structural limitations, ligand chemistry structure and drug discovery project decisions. Adv Drug Deliv Rev 101, 34–41, doi:10.1016/j.addr.2016.04.029 (2016).

21 Price, E. et al. Beyond Rule of Five and PROTACs in Modern Drug Discovery: Polarity Reducers, Chameleonicity, and the Evolving Physicochemical Landscape. J Med Chem 67, 5683–5698, doi:10.1021/acs.jmedchem.3c02332 (2024).

22 Furukawa, A. et al. Drug-Like Properties in Macrocycles above MW 1000: Backbone Rigidity versus Side-Chain Lipophilicity. Angew Chem Int Ed Engl 59, 21571–21577, doi:10.1002/anie.202004550 (2020).

23 Johns, D. G. et al. Orally Bioavailable Macrocyclic Peptide That Inhibits Binding of PCSK9 to the Low Density Lipoprotein Receptor. Circulation 148, 144–158, doi:10.1161/CIRCULATIONAHA.122.063372 (2023).

24 Tanada, M. et al. Development of Orally Bioavailable Peptides Targeting an Intracellular Protein: From a Hit to a Clinical KRAS Inhibitor. J Am Chem Soc 145, 16610–16620, doi:10.1021/jacs.3c03886 (2023).

25 Klein, V. G., Bond, A. G., Craigon, C., Lokey, R. S. & Ciulli, A. Amide-to-Ester Substitution as a Strategy for Optimizing PROTAC Permeability and Cellular Activity. J Med Chem 64, 18082–18101, doi:10.1021/acs.jmedchem.1c01496 (2021).

26 Hosono, Y. et al. Amide-to-ester substitution as a stable alternative to N-methylation for increasing membrane permeability in cyclic peptides. Nat Commun 14, 1416, doi:10.1038/s41467-023-36978-z (2023).

27 Taechalertpaisarn, J., Ono, S., Okada, O., Johnstone, T. C. & Lokey, R. S. A New Amino Acid for Improving Permeability and Solubility in Macrocyclic Peptides through Side Chain-to-Backbone Hydrogen Bonding. J Med Chem 65, 5072–5084, doi:10.1021/acs.jmedchem.2c00010 (2022).

28 Gibson, W. J. et al. Bifunctional Small Molecules That Induce Nuclear Localization and Targeted Transcriptional Regulation. J Am Chem Soc 145, 26028–26037, doi:10.1021/jacs.3c06179 (2023).

29 Simpson, L. M. et al. Target protein localization and its impact on PROTAC-mediated degradation. Cell Chem Biol 29, 1482–1504 e1487, doi:10.1016/j.chembiol.2022.08.004 (2022).

30 Deshaies, R. J. Multispecific drugs herald a new era of biopharmaceutical innovation. Nature 580, 329–338, doi:10.1038/s41586-020-2168-1 (2020).

